# Life-history traits explain variation in the larval dispersal potential of corals

**DOI:** 10.64898/2026.09.29.755390

**Authors:** Matthew J Morecroft, Ariel Greiner, Lena L Faber, Katrina J Davis

## Abstract

1. Protecting and restoring corals is critical to conserve marine biodiversity and ecosystem services in the Anthropocene. Knowledge of coral larval dispersal dynamics can be used to identify important sites for coral metapopulation connectivity and to spatially optimise coral conservation efforts. However, we know little regarding the larval dispersal of many coral species and our understanding of the relationships between coral larval dispersal and life-history traits (e.g., reproductive mode, colony morphology), phylogenetic relationships, and biogeography is incomplete. These knowledge gaps impede coral conservation.
2. In this research, we systematically reviewed the species, regions, and methodologies of studies of coral larval dispersal. Additionally, we conducted a meta-analysis to quantify the heterogeneity in reported coral larval duration values and identified the roles of life-history traits in driving that heterogeneity.
3. We found that various methods (experimental, biophysical modelling, and population genetics) were used to study 221 coral species across many regions. Soft and temperate corals received comparatively lower research attention. Our meta-analysis reveals that reproductive mode and colony morphology explain significant heterogeneity in larval settlement times, and that closely-related species have more similar larval settlement times.
4. Our findings suggest that reproductive mode and colony morphology could be used to predict larval duration in under-studied coral species, and that larval duration could be approximated from closely-related corals when trait data is unavailable. Our results support strategies focused on modelling differing larval durations across corals with different life-history traits to predict realised dispersal in under-studied coral species and identify conservation priority sites.

## Introduction

Effectively protecting corals from anthropogenic threats is an important conservation priority, owing to their importance to marine ecosystems and the high levels of threats they face. Corals have critical roles as habitat builders across many marine environments and support high levels of marine biodiversity and ecosystem services (Cordes et al., 2023; Fisher et al., 2015). Many corals are highly vulnerable to the impacts of climate change, due to rapidly warming shallow waters and susceptibility to bleaching at high temperatures (IPCC, 2022). Intensive fishing, pollution, and other anthropogenic stressors also threaten a wide variety of coral populations (Cordes et al., 2023; Gove et al., 2023). The persistence of coral populations in the face of such threats depends on several factors, including potential for spatial rescue (Greiner et al., 2022), gene flow for adaptation (Vogt-Vincent et al., 2023), and potential range expansion (Vogt-Vincent et al., 2025). These factors all rely on sufficient coral larval dispersal connecting coral metapopulation patches to facilitate reef recovery and resilience. Connectivity is therefore an important consideration in the designation of marine protected areas and coral restoration projects, which are most effective when focused on reefs that are important larval sources or “stepping stones” for metapopulation connectivity (Brookson and Greiner, 2026; Greiner et al., 2026; King et al., 2023). A lack of data on the larval dispersal dynamics and connectivity of many coral populations impedes effective conservation measures (Burt et al., 2024; Randall et al., 2024; Vogt-Vincent et al., 2024).

Connectivity in coral metapopulations is mainly driven by passive dispersal during the motile planula larval stage (Gleason and Hofmann, 2011). Determining coral population connectivity requires knowledge of corals’ realised and/or potential larval dispersal. Realised dispersal refers to the actual distances travelled by larvae in the field from release to successful settlement on appropriate substrate. Dispersal potential is a property describing the theoretical travel capabilities of larvae. Realised larval dispersal is largely passive and depends on various oceanographic factors, including: currents, temperature, settlement substrates, chemical cues, light, sound, and sedimentation (Gleason and Hofmann, 2011; Vogt-Vincent et al., 2023). Additionally, realised dispersal is dependent on dispersal potential. Dispersal potential is driven by factors including: larval duration (i.e., time taken for a larva to either settle or die), swimming/crawling behaviour, larval weight, and timing of larval release (Connolly and Baird, 2010; Strader et al., 2018). Larval duration can be affected by mortality, development of settlement competency, and settlement initiation behaviours (Aoki et al., 2024; Harrison and Wallace, 1990). Realised dispersal is difficult to directly measure in situ due to small larval sizes, large quantities, infrequent production, and inability to track larvae (Vogt-Vincent et al., 2023). Technologies, notably biophysical modelling and genetic sequencing, have developed over time to overcome these challenges and enabled increased research attention towards realised larval dispersal (Alvarado-Cerón et al., 2023).

Several biological factors may explain variation in coral dispersal potential, including reproductive mode, colony morphology, and biogeography. The most well-understood life-history trait influencing larval dispersal potential is reproductive mode, describing whether planulae are “broadcasted” (fertilised externally) or “brooded” (fertilised internally). Researchers have observed several brooding corals to have shorter larval durations or smaller-scale connectivity patterns compared to some broadcasting species (Davies et al., 2017; Ritson-Williams et al., 2009; van der Ven et al., 2021). However, differences in dispersal potential between the two reproductive modes have not been systematically tested across a large number of species from across biogeographic realms. Previous analysis of coral life-history strategies has found that reproductive mode, colony morphology, and growth rate were the three most important life-history traits differentiating coral life-history strategies, and that biogeographic realm influenced the relative importance of different traits (Darling et al., 2012). Different coral life-history strategies, including competitive, weedy, stress-tolerant and generalist strategies, are associated with different levels of investment into survival, growth, and reproduction (Darling et al., 2012; Rachello-Dolmen and Cleary, 2007). As different life-history strategies invest differently into larval production and may benefit differently from varying levels of local larval retention, colony morphology, growth rate, and biogeographic realm could also correlate with dispersal potential (Darling et al., 2012; Otis et al., 2024; van Woesik et al., 2012). Life-history strategies are often phylogenetically conserved, and so coral larval dispersal potential may also be influenced by phylogeny. However, the extent to which colony morphology, growth rate, biogeographic realm, and phylogeny correlate with dispersal potential has not been evaluated. Therefore, determining the ability of life-history traits and biogeographic realm to explain variation in dispersal potential could indicate which traits could more reliably predict realised dispersal in data-poor coral populations.

Different methods used to estimate realised or potential coral larval dispersal have various limitations. Population genetics (and genomics) approaches indirectly measure realised dispersal in situ. Such methods are often costly and can sometimes be limited by confounding effects of post-settlement processes (Alvarado-Cerón et al., 2023; Burt et al., 2024; Macleod et al., 2024). Biophysical models are lower-cost, combining oceanographic information with biological information on dispersal potential to predict realised dispersal patterns (Vogt-Vincent et al., 2023). Modelling can capture variability in dispersal distances, but can be limited by the quality of available information (Figueiredo et al., 2022; Vogt-Vincent et al., 2024). Empirical data from experimental studies measuring larval dispersal potential can be used to parameterise biophysical models, although many models are parameterised on data from different coral species than the population being modelled (Macleod et al., 2024). Experimental studies are mostly laboratory-based, with very few in-situ studies of realised or potential dispersal (Sammarco and Andrews, 1988; Suzuki et al., 2011), and estimate coral dispersal potential by measuring either survival-or settlement-based metrics of larval duration (Randall et al., 2024). It is currently unclear how consistent coral larval duration estimates are between experimental studies, these values can vary widely even within a single species (e.g., mean larval settlement time in *Acropora tenuis* differs by at least 40 days across studies) (Moneghetti et al., 2019; Suzuki et al., 2011). Addressing knowledge gaps around the extent and drivers of variability in experimental larval duration estimates will therefore enhance both our understanding of coral larval dispersal potential and models predicting realised dispersal.

In this study, we assess coral larval duration (a key determinant of coral larval dispersal potential) and its key drivers using meta-analysis and identify species and regional knowledge gaps in larval dispersal using a systematic review. We identified relevant studies of coral larval dispersal for the systematic review using the Web of Science and Scopus databases. For each study included in the systematic review, we determined the methods authors used, and the species and biogeographic realms of the coral populations they studied. To quantify heterogeneity in reported coral larval durations, we extracted data describing either larval settlement or larval survival times available from experimental studies to use in a random-effects meta-analysis. Using mixed-effects meta-regressions and publicly available data on reproductive mode, colony morphology, phylogeny, and biogeographic realm, we tested the effects of these biological factors on heterogeneity in larval durations. This study assesses the taxonomic and regional extent of our knowledge of coral larval dispersal, a historically under-studied aspect of coral biology but one that is crucial to understand when designing management plans for coral reefs.

## Methods

### Replication Statement

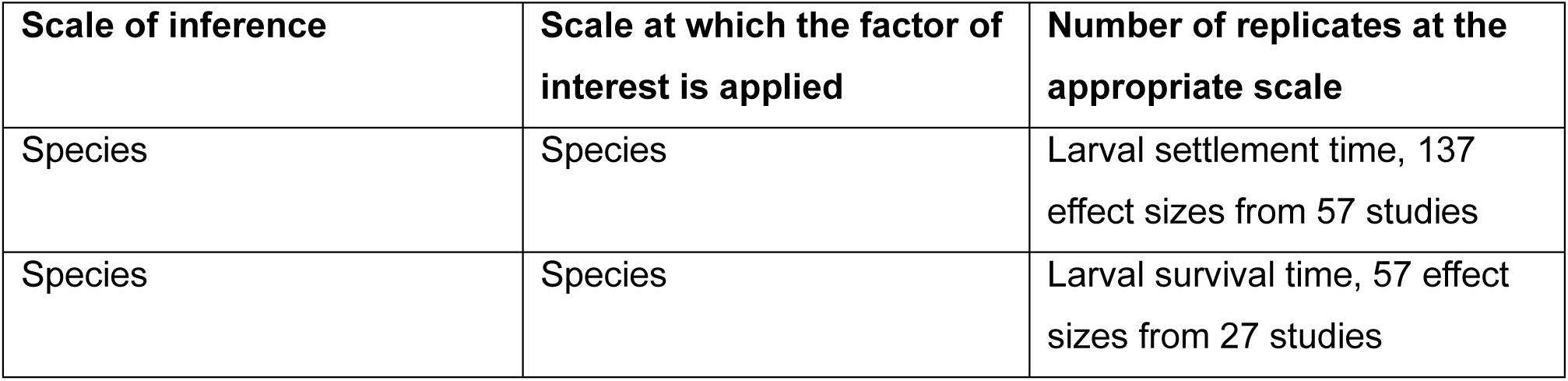

| Scale of inference | Scale at which the factor of interest is applied | Number of replicates at the appropriate scale |
| --- | --- | --- |
| Species | Species | Larval settlement time, 137 effect sizes from 57 studies |
| Species | Species | Larval survival time, 57 effect sizes from 27 studies |

### Overview

To assess variation in reported coral larval durations and examine the effects of life-history traits on this variation, we conducted a meta-analysis of coral larval duration (specifically, coral larval settlement time and coral larval survival time). To identify species, regions, and methods used in studies of coral larval dispersal, we conducted a systematic review of the coral larval dispersal literature. We identified relevant scientific literature from the Web of Science and Scopus databases and extracted data on methodologies, study taxa, and study regions from publications included in our review. To obtain meta-analysis effect sizes, we extracted mean larval settlement times and mean larval survival times from experimental studies with sufficient available data. To evaluate the extent of heterogeneity in effect sizes, we used random-effects meta-analysis models. To evaluate the contribution of life-history traits to effect size heterogeneity, we constructed several mixed-effects meta-regression models. We obtained trait and phylogeny data from the Coral Trait Database and the Open Tree of Life respectively.

### Systematic review

We used the Web of Science and Scopus databases to search widely across the coral larval dispersal literature (Koricheva et al., 2013). To identify publications relevant to coral larval dispersal and ensure search reproducibility, we applied a 3-part search string to similar fields over a similar timeframe in both databases (**Supporting Methods - 1**). Our search produced 6,192 results, with 4,287 unique publications (**Figure S1**). To ensure only relevant studies were included in the systematic review, we manually screened the identified studies’ titles and abstracts for relevance, leading to the exclusion of 3,287 studies (**Figure S1**). Studies with the full text unavailable (n= 34) were also removed. We assessed whether the remaining studies were eligible for inclusion in the systematic review, using three inclusion criteria. Firstly, studies had to contain primary research. Secondly, studies had to be of extant coral populations. As definitions of coral can vary, we defined corals here as anthozoans, excluding orders Actiniaria (true anemones) and Ceriantharia (tube-dwelling anemones). This ensured that all corals in this study had comparable larval characteristics (Coelho and Lasker, 2016; Stampar et al., 2015). Thirdly, studies had to research processes relevant to larval dispersal, including population connectivity and recruitment. After screening, 653 studies were included in the systematic review (**Figure S1; Table S1**).

### Data extraction

To compare measured coral larval durations between experimental studies using meta-analysis, we used two effect sizes: mean larval settlement time and mean larval survival time. The mean larval settlement time refers to the mean time at which a larva settled in a specific study; the mean survival time refers to the mean lifespan (i.e., the mean time of mortality) of a larva in a study. We used these measures as larval mortality is considered a separate property to larval settlement competency and is typically experimentally measured in the absence of settlement cues and so the two effect sizes cannot be directly compared (Connolly and Baird, 2010). Both times are typically measured in days. Effect sizes were recorded alongside the variance of their estimated means, calculated from squaring the standard error (SE) in larval settlement/survival times. We extracted effect sizes from all experimental studies (both lab-based and field-based) where they were available, using several extraction protocols (**Supporting Methods - 2**).

Testing the effects of different biological factors on coral larval durations required data on coral life-history traits and locations from studies reporting effect sizes (n= 64). We extracted study location from the latitude and longitude of the midpoint of the area studied. Comprehensive ordination approaches to integrate multiple traits and categorise life-history strategies (*sensu* Darling et al. (2012)) have only been conducted on a limited range of coral species in our meta-analysis. However, such analyses identified reproductive mode, colony morphology, and growth rate as the three most important traits differentiating coral life-history strategies (Darling et al., 2012). Data on reproductive mode and colony morphology were available for a much wider range of species (Gómez-Gras et al., 2025; Madin et al., 2016). Other traits, including growth rate, were only available for a small number of species in this analysis. Therefore, we focused on reproductive mode and colony morphology as the relevant life-history trait variables in our meta-analysis. The reproductive mode of each species was obtained from each paper’s main text and recorded as either broadcasting or brooding. We downloaded additional data on colony morphology from the Coral Trait Database (Madin et al., 2016) and categorised coral species into 5 different morphological groups (**Supporting Methods - 3; Figure S2; Figure S3**).

To review trends in coral larval dispersal research, we extracted data on study methodology from all eligible studies (n= 653). We determined what categories of methodology were used in each study to estimate dispersal. Categories included experimental, biophysical modelling, and population genetics methods, with some studies falling into multiple categories. We classified lab-based and field-based methods as sub-categories of experimental approaches. We also recorded the species (or other taxonomic grouping) and marine biogeographic realm studied (according to Spalding et al. (2007)). To obtain detailed taxonomic information for each species studied, we downloaded anthozoan taxonomic data from the World Register of Marine Species (WoRMS Editorial Board, 2026). We used data from the IUCN Red List to identify the conservation status of coral species in this review (IUCN, 2025). To assess how empirical larval duration estimates are used in biophysical models of larval dispersal potential, we recorded the justifications given for larval duration parameters used in biophysical modelling studies. We also recorded whether empirical data from corals of the same species, genus, family, order, or reproductive mode as the study species were used as a source for these model parameters.

### Meta-analysis

To determine the level of heterogeneity in estimates of coral larval duration, random-effects meta-analysis models were constructed for both mean larval settlement time and mean larval survival time. All statistical analyses were conducted in R (v4.6.1) (R Core Team, 2024). To account for the different levels of precision in each study’s estimate of effect size, all the meta-analysis models we constructed weighted each effect size by the inverse of its variance (Koricheva et al., 2013). For the first stage of the meta-analysis, we constructed random effects models to quantify the overall heterogeneity in effect sizes, using the “*metafor*” package (v 5.2-1) with a restricted maximum likelihood estimation (REML) method (Koricheva et al., 2013; Viechtbauer, 2010). To account for effect sizes reported from the same study being more similar to each other than those reported from other studies, these models contained a random between-observation variance component nested within a random between-study variance component. This meant effect sizes varied around mean values for a study, which in turn varied around an overall mean (Koricheva et al., 2013). After model construction we calculated: the σ^2^ value for each random-effects variance component, the effect size heterogeneity (Q) (i.e., variation in effect sizes between studies relative to that expected from sampling error), and Akaike information criterion (AIC) values.

As species with shared evolutionary history are more likely to have similar traits, it is important for meta-analysis models to account for potential phylogenetic non-independence of effect sizes (Koricheva et al., 2013; Lajeunesse, 2009). Therefore, we constructed phylogenetically-controlled random-effects meta-analysis models for both mean larval settlement and mean survival time effect sizes. These models contained a variance component describing between-species variation. To define how closely coral species were related to each other, we linked this variance component to a phylogenetic correlation matrix (Stotz et al., 2025). This matrix was generated from a phylogeny from the Open Tree of Life Synthetic Tree (OpenTreeOfLife et al., 2019) using the “*rotl*” (v3.1.1) and “*ape*” packages (v5.8-1) (**Fig S2**) (Michonneau et al., 2016; Paradis and Schliep, 2019). To determine whether phylogenetic relationships had a meaningful role in explaining variation in effect sizes, we assessed phylogenetically-controlled model performance using AIC values and compared to the performance of non-phylogenetic random-effects models (Koricheva et al., 2013; Lajeunesse, 2009).

To test whether life-history traits and biogeographic variation were significant drivers of heterogeneity in coral larval durations, we used mixed-effects meta-regression models. We also constructed these models with “*metafor*”, using a REML method (Viechtbauer, 2010). For both effect sizes (mean settlement and survival times), each model contained different fixed-effect moderator variables. These variables were: reproductive mode, colony morphology, temperate vs. tropical region, marine biogeographic realm, and latitude. To examine how different life-history traits interacted in their effects on larval duration, reproductive mode and colony morphology with an interaction effect were also included as fixed effects in the same model. This ensured that the models accounted for potential differences in the role functional traits play in different life-history strategies, as reproductive mode has been shown to co-vary with colony morphology across life-history strategies (Darling et al., 2012; Rachello-Dolmen and Cleary, 2007). Each mixed-effects model also contained the random-effects variance components from the random-effects models, with phylogenetic control included if the phylogenetically-controlled model was the best-fitting random-effects model (Stotz et al., 2025). We calculated the heterogeneity in effect sizes attributed to the moderator variable for each model, and tested this value for significance using the “*metafor*” omnibus test of moderators. The residual heterogeneity was also calculated and tested for significance using “*metafor*”’s Q_E_ test (Koricheva et al., 2013; Viechtbauer, 2010). When moderator variables were significant, we reported the model estimates of mean duration for specific groups of corals.

## Results

### Systematic review

Our systematic review showed that the number of studies of coral larval dispersal published each year has increased over time (**Figure 1a**). We identify 653 studies with primary research on coral larval dispersal. Of these, 311 use experimental methods (with 213 studies using lab-based and 123 field-based methods), 237 use population genetics, and 184 use biophysical modelling, with several studies (n= 73) combining different methodologies. Studies published per year have increased within each study methodology category (**Figure 1a**). Of the experimental studies, 64 have sufficient data available to extract meta-analysis effect sizes.

**Figure 1:**
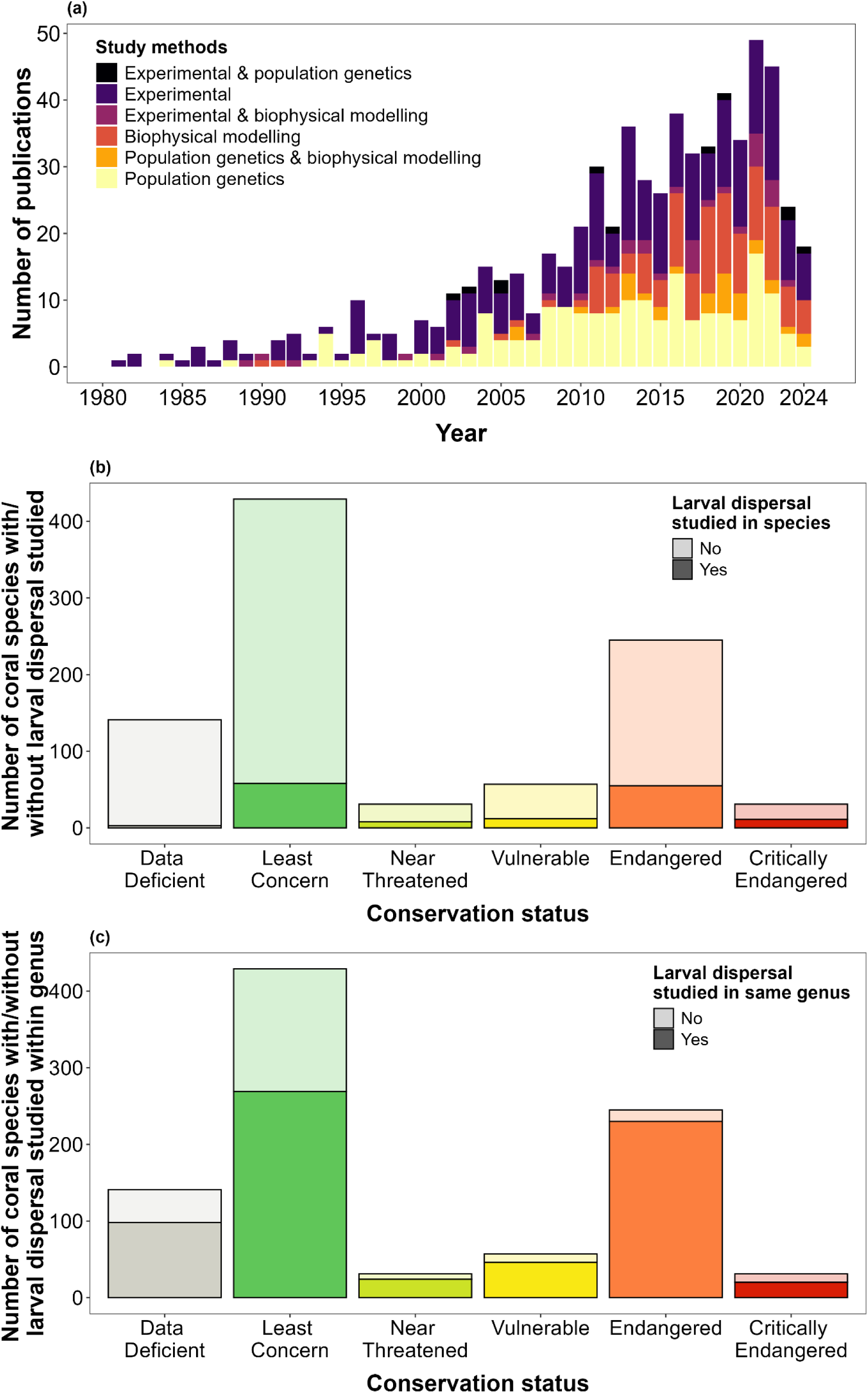
Methods and species included in studies of coral larval dispersal, 1980 – 2024. (a) Bar plots showing the methodologies in studies of coral larval dispersal published from 1980 to 2024. Study totals are broken down by methodology over time. Studies identified from Web of Science only include those published prior to June 2024 and studies from Scopus only go up to January 2024, hence the shorter 2024 bar. (b & c) The number of coral species included in these studies are broken down by IUCN Red List status (IUCN, 2025). Bold colours indicate species included in a study (b) or where at least one species from a genus is included in these studies (c). Note that of all currently accepted coral species, 5,035 have not been evaluated by the IUCN Red List (n= 5,035), so are not shown.

The coral larval dispersal literature has focused on a variety of corals, including 221 species from 115 genera. Of these, 158 species across 69 genera were stony corals (representing 9.3% of the order Scleractinia), and 63 species across 46 genera were soft corals (1.5% of named non-Scleractinian corals). Larval dispersal has not been studied in 76.6% of coral species classified as Threatened under the IUCN Red List (IUCN, 2025) (**Figure 1b**). However, genus-level coverage was much greater, especially among Endangered and Data Deficient species (**Figure 1c**).

The areas studied in the literature covered a wide range of latitudes (67°S-70°N) and longitudes (180°W-180°E). The most widely studied biogeographic realms were the Central Indo-Pacific (n= 265), the Tropical Atlantic (n= 156), and the Temperate Northern Atlantic (n= 99). There were 518 studies of tropical, 179 of temperate, and 8 of polar populations, reflecting a research focus towards tropical over cold-water corals. Notably, despite large contributions of Indian Ocean populations to stony (Kusumoto et al., 2020) and soft coral diversity (Pérez et al., 2016), relatively few studies research coral larval dispersal in the Western Indo-Pacific and Temperate Southern Africa (**Figure 2**).

**Figure 2:**
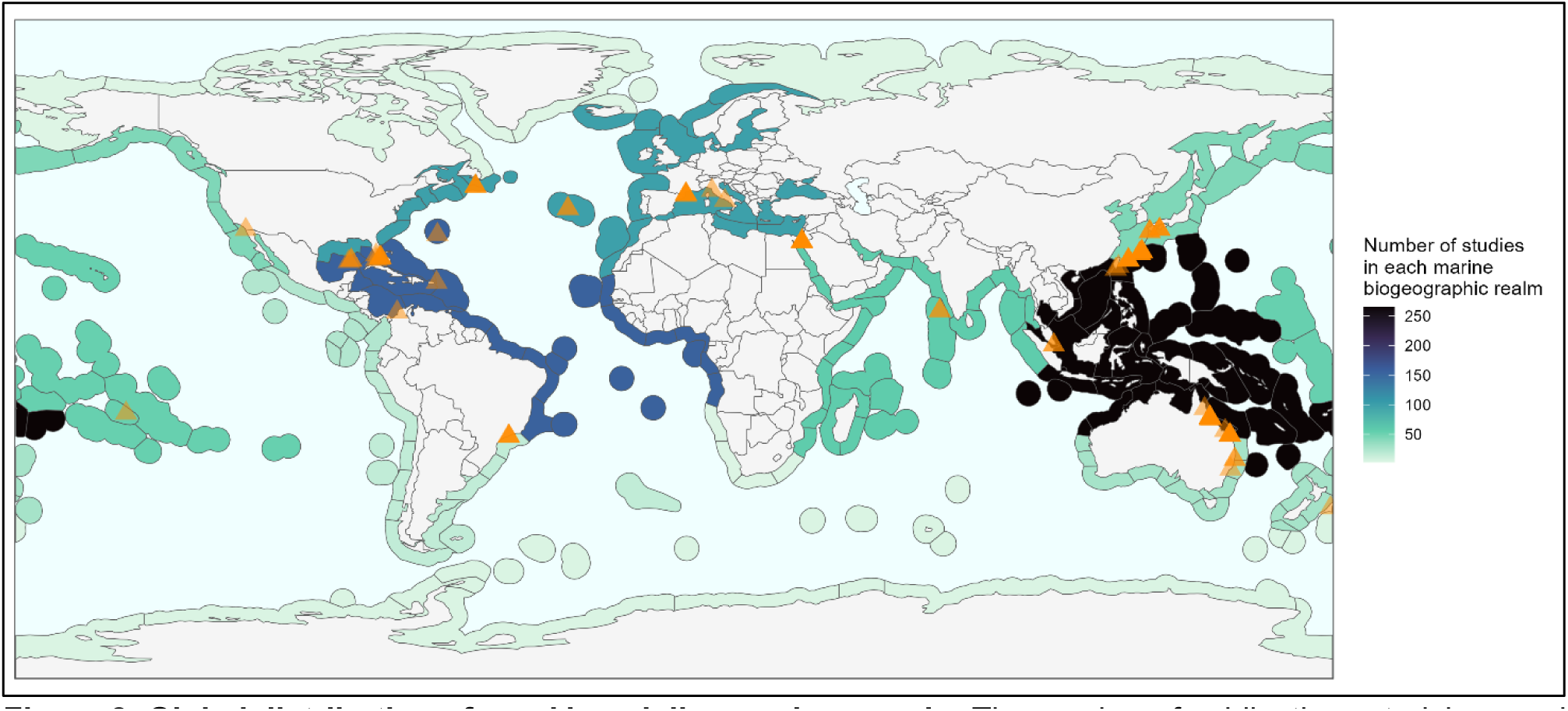
Global distribution of coral larval dispersal research. The number of publications studying coral larval dispersal in each marine biogeographic realm (Free, 2026; Spalding et al., 2007). Colour indicates the number of studies in each realm. Triangles indicate locations of experimental studies included in our meta-analysis.

Of 184 biophysical modelling studies, 166 use model parameters relating to larval duration, including: larval competency gain/loss rates, mortality rates, and maximum pelagic larval duration (PLD) values. There are 29 studies modelling corals with a range of larval duration parameters for reasons including: species’ larval duration was unknown; study assessed the effects of varying larval duration on dispersal distances; models were generalised for a variety of species. Of 137 remaining studies, 102 use empirical data to justify the larval duration parameter value, with the other 35 providing no clear justification for this parameter. Of these 102 studies, 11 collect their own experimental data to parameterise larval duration, while the remaining 91 studies use data from previous literature. Notably, 42 of these studies (41.2%) cite the same publication (Connolly and Baird, 2010), with this being the only study cited to justify the choice of larval duration parameter in 27 cases. Not all models are parameterised using data from the same study species (**Figure 3**). Generic corals from different groups are modelled in 82 studies (e.g., “generic stony coral”, “generic Acropora”, “generic broadcaster” etc.), which can only cite empirical data with limited specificity.

**Figure 3:**
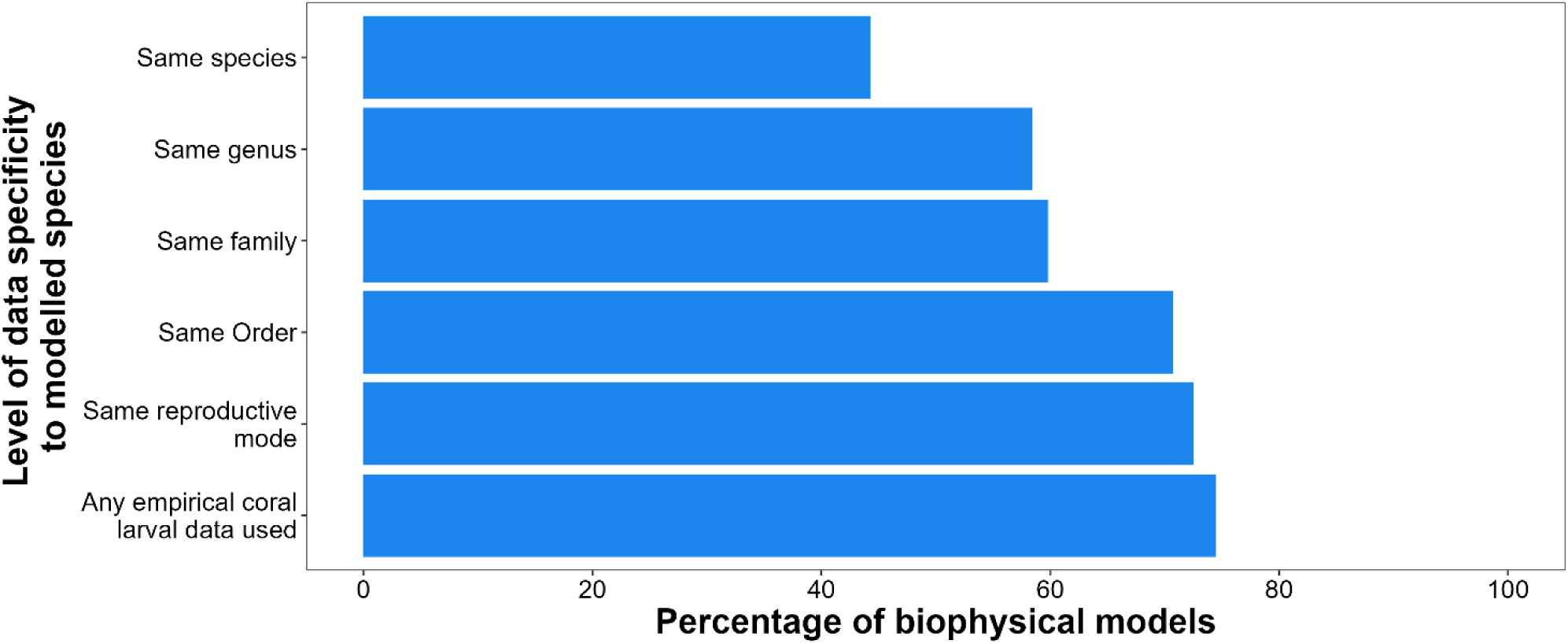
Specificity of empirical data used to parameterise larval duration in biophysical models of coral larval dispersal. The percentage of biophysical model larval duration parameters that cite empirical data from the same species, genus, family, order, or reproductive mode, as that being studied, or from any coral. Each percentage is calculated only for models containing a specific larval duration parameter for specific corals at each level (n= 88, 101, 102, 123, 131, 137, descending from species to coral) (i.e., excludes models of generic corals from the total for each level, and excludes models using a range of larval duration parameters).

### Meta-analysis

We assess the extent of variation in reported coral larval durations using random-effects meta-analysis models, finding a significant level of heterogeneity in effect sizes. The best-fitting random-effects models predict an overall mean larval settlement time of 15.6 ±3.5 days (±SE) and a mean survival time of 25.4 ±3.9 days, with heterogeneity values of Q= 30,5013.2 and Q= 146,345.7 respectively (**Table 1**). The heterogeneity (Q) determined by the random-effects models is significant in all models (p<0.0001). This indicates that variation between measurements of effect sizes was significantly greater than variation expected due to sampling error alone, showing that estimated coral larval durations vary significantly between observations in the literature. The phylogenetically-controlled models provide the best fits for mean larval settlement time, but not for mean larval survival time (**Table 1**).

**Table 1:** Table summarising the outputs (AIC, Heterogeneity, and p-value obtained from Cochran’s Q-test of heterogeneity) from random-effects meta-analysis models of effect sizes for both mean larval settlement time and mean larval survival time. Models with “species” as a random effect refer to models which control for phylogeny.

| Random effects | Mean larval settlement time |  |  | Mean larval survival time |  |  |
| --- | --- | --- | --- | --- | --- | --- |
|  | AIC | Heterogeneity | Test for | AIC | Heterogeneity | Test for |
|  |  | (Q) | heterogeneity<br>(p-value) |  | (Q) | heterogeneity<br>(p-value) |
| study \| observation | 1058.6 | 340922.0 | <0.0001 | 523.3 | 146345.7 | <0.0001 |
| study \| observation<br>+ species | 1053.4 | 340922.0 | <0.0001 | 524.2 | 146345.7 | <0.0001 |

Mean larval settlement time varies significantly with reproductive mode and colony morphology. The meta-regression model of mean larval settlement time with reproductive mode and colony morphology as fixed-effects moderator variables has a moderator heterogeneity value of Q_M_= 40.149, which the Q_M_-test determined was significant (p<0.0001, **Table 2**). This indicates that reproductive mode and colony morphology explain significantly more variation in mean larval settlement times than expected from sampling error alone. Comparing random-effects σ^2^ values between this model and the corresponding phylogenetically-controlled random-effects model indicates that nearly all the variation attributed to phylogeny in the random-effects model was encompassed by colony morphology and reproductive mode (**Table S4**). Differences between mean larval settlement times of broadcast and brooded larvae varied across different colony morphologies (**Figure 4a; Table S5**). The corresponding meta-regression for mean larval survival time found that reproductive mode and colony morphology were not significant predictors (Q_M_= 7.804; p= 0.453) (**Figure 4b**). For all mixed-effects meta-regression models, the residual heterogeneity (Q_E_) is significant (**Table 2**).

**Figure 4:**
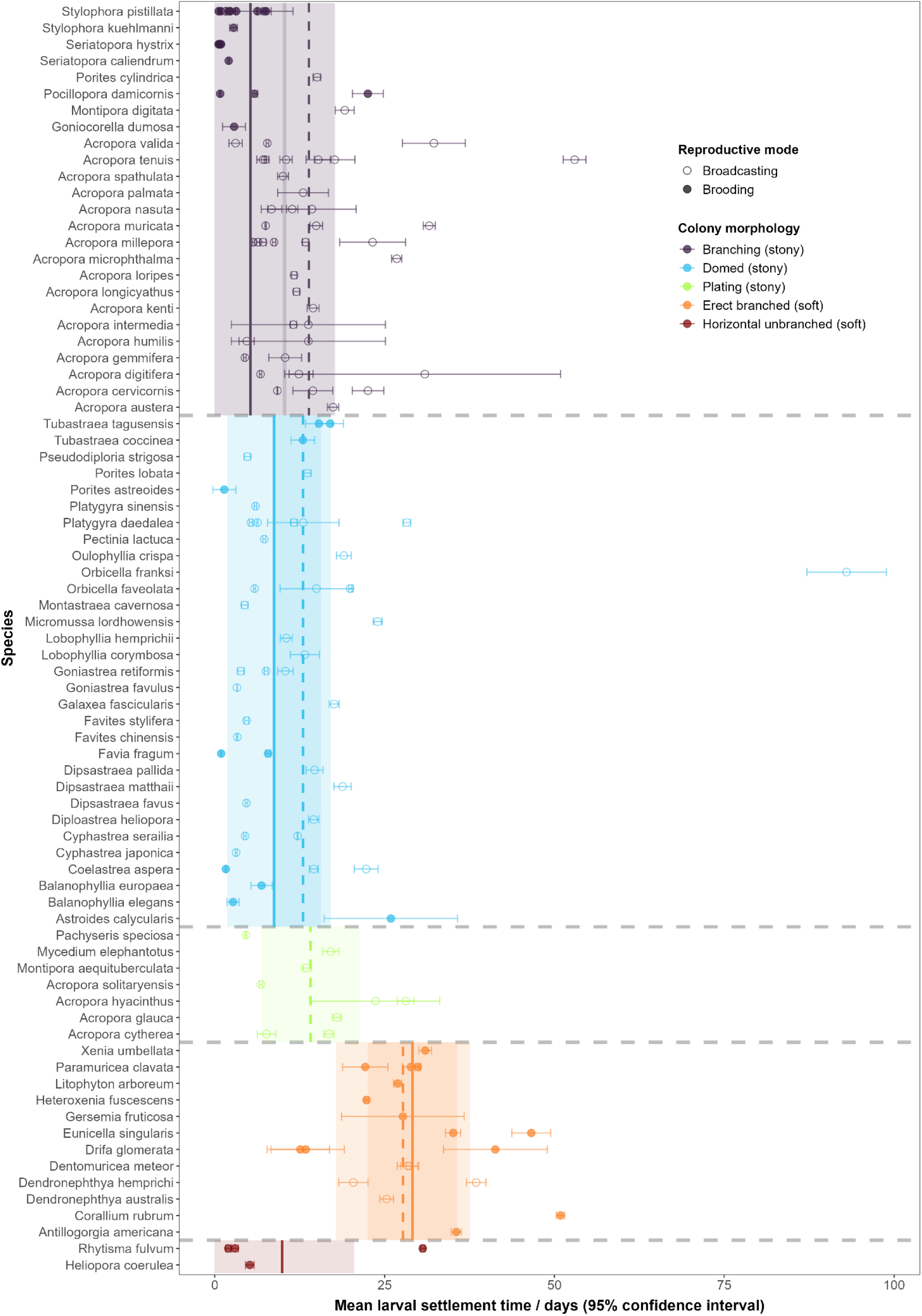
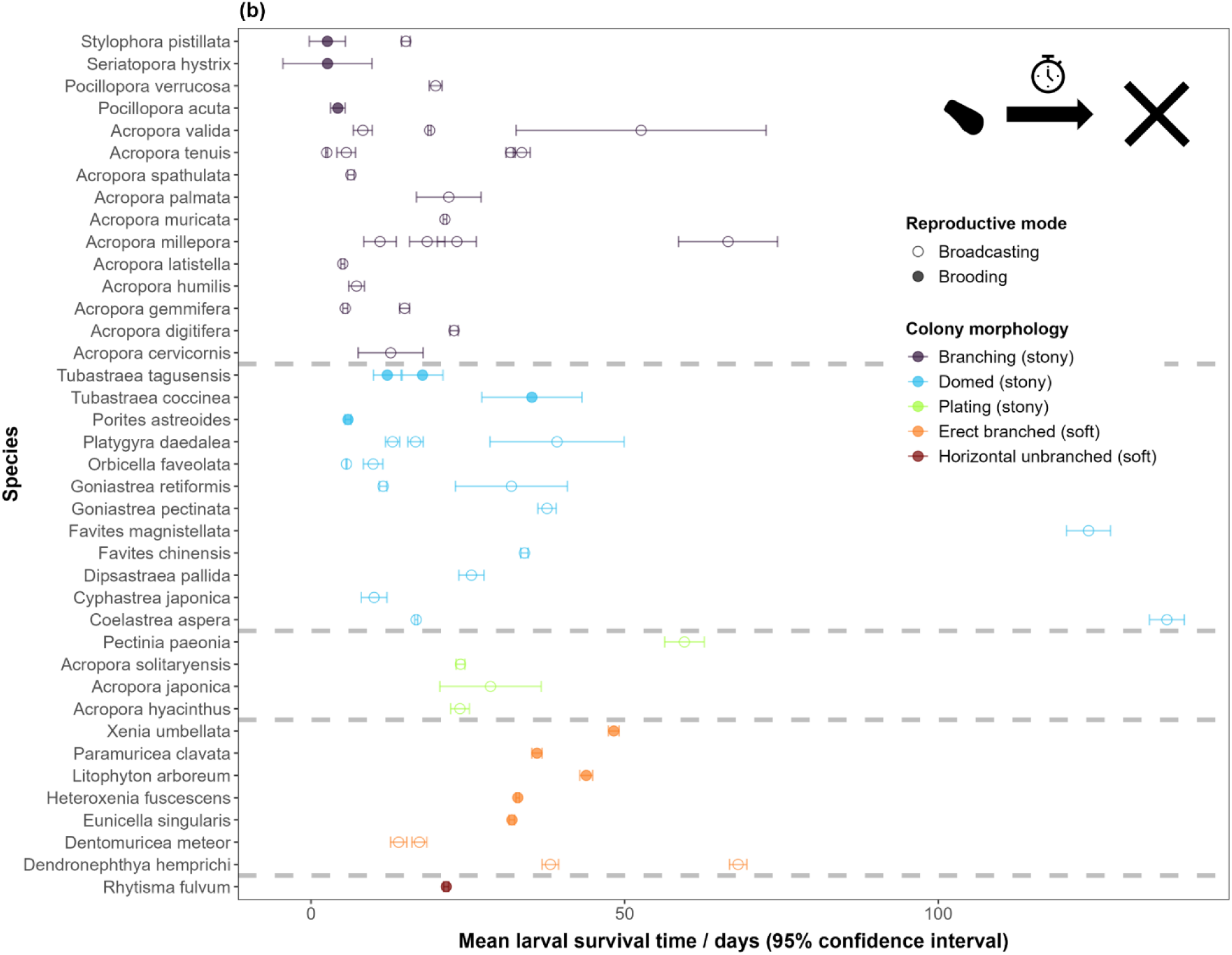
Life-history traits explain variation in mean larval settlement time, but not in mean larval survival time. Forest plots showing variation in effect sizes for mean larval settlement time (a) and mean larval survival time (b) across coral species, colony morphologies, and reproductive modes. Points with error bars represent individual effect sizes with 95% confidence interval. Vertical lines represent overall mean larval settlement time estimated for broadcasting (dashed) or brooding (solid) corals for each colony morphology. Shaded area indicates 95% confidence interval. Reproductive mode and colony morphology did not significantly contribute to heterogeneity in mean larval survival time, so no vertical lines used in (b).

**Table 2:** Table summarising outputs (Moderator heterogeneity, p-value obtained from QM-test of moderator heterogeneity, Residual heterogeneity, and p-value obtained from QE-test of residual heterogeneity) from mixed-effects meta-regression models of effect sizes for both mean larval settlement and mean larval survival times. Reproductive mode*Colony Morphology refers to the interaction effect between the two life-history trait variables used in their corresponding meta-regression.

| Effect size | Moderators | Moderator<br>heterogeneity<br>(Q<sub>M</sub>) | Test of<br>moderator<br>(p-value) | Residual<br>heterogeneity<br>(Q<sub>E</sub>) | Test of residual<br>heterogeneity<br>(p-value) |
| --- | --- | --- | --- | --- | --- |
| Mean larval<br>settlement time | Reproductive mode*<br>Colony morphology | 39.953 | <0.0001 | 203185.1 | <0.0001 |
|  | Tropical/temperate<br>region | 1.338 | 0.247 | 262496.2 | <0.0001 |
|  | Marine<br>biogeographic realm | 8.435 | 0.296 | 186649.8 | <0.0001 |
|  | Latitude | 0.775 | 0.379 | 335475.7 | <0.0001 |
| Mean larval<br>survival time | Reproductive mode*<br>Colony morphology | 7.611 | 0.368 | 68948.8 | <0.0001 |
|  | Tropical/temperate<br>region | 0.008 | 0.930 | 128171.2 | <0.0001 |
|  | Marine<br>biogeographic realm | 4.339 | 0.631 | 60257.6 | <0.0001 |
|  | Latitude | 0.183 | 0.669 | 114436.4 | <0.0001 |

Biogeographic variation does not have a significant influence on coral larval durations. Mixed effects meta-regressions were fitted using temperate/tropical region, marine biogeographic realm, and study median latitude as moderator variables. For both larval settlement and survival time, neither temperate/tropical zone, marine biogeographic realm, nor latitude contribute to a significant level of heterogeneity in effect sizes (**Table 2**).

## Discussion

In this research, we find that life-history traits drive variation in coral larval dispersal potential. Estimates of coral larval durations are highly variable across the published literature, and our analyses show that reproductive mode and colony morphology are significant moderators of heterogeneity in mean larval settlement time. Controlling for phylogeny provides better-fitting random-effects meta-analysis models of settlement time compared to models without these controls, but phylogenetic relationships did not contribute any variation in larval settlement time not already explained by reproductive mode and colony morphology. Our systematic review shows that larval dispersal has been assessed using various methods in 221 coral species from a variety of biogeographic realms. However, some regions and species receive disproportionately less attention, and the larval dispersal of many coral species (5,738) have not been studied in the published literature. Although the majority of biophysical models use empirical data to inform larval duration parameter values, we find that only 44.3% of empirically-based models of a named species were parameterised using data from the same species (**Figure 3**). However, our results indicate that available life-history trait data and closely-related species’ larval duration data could be used to predict dispersal potential effectively in coral species where larval dispersal remains under-studied.

Meta-analyses of coral larval durations reveal that variation in larval settlement time can be explained in part by the reproductive mode and morphology of corals. Our meta-regression testing the contributions of reproductive mode and colony morphology towards variation in larval settlement time shows these variables to be significant (**Table 2**), suggesting that these life-history traits are important drivers of coral larval dispersal potential. This finding aligns with some previous observations of brooding corals with smaller-scale connectivity patterns compared to broadcasters in the same area (Davies et al., 2017; Ritson-Williams et al., 2009; van der Ven et al., 2021). Faster settlement in brooded larvae could potentially be explained by their increased development upon release (Harrison and Wallace, 1990; Kahng et al., 2011). Our finding that colony morphology drives variation in larval settlement times is noteworthy, given the apparent paucity of research explicitly linking colony morphology to connectivity dynamics in corals. Additionally, our findings align with research highlighting the contribution of colony morphology to life-history strategies and functional roles in stony and soft corals (Darling et al., 2012; Otis et al., 2024; Velásquez and Sánchez, 2015). Differing investments in reproduction between corals with different morphologies could be responsible for driving differences in larval duration. Several studies have shown that reproductive traits co-vary with adult growth traits in corals; for instance, stress-tolerant stony corals associate with both domed morphologies (influencing light and heat exposure) and broadcasted larvae (produced in large numbers with reduced investment in each larvae) (Darling et al., 2012; van Woesik et al., 2012). Our findings suggest that information on reproductive mode and colony morphology could be used to estimate larval settlement times, and consequent patterns of dispersal and connectivity, in coral species without any larval dispersal potential information.

Results from the phylogenetically-controlled meta-analysis models suggest that higher-taxa information may also be useful in addressing coral larval dispersal knowledge gaps. Accounting for phylogeny provides better-fitting random-effects models of mean larval settlement times (**Table 1**), indicating that more closely related species were more likely to have similar larval settlement times (Lajeunesse, 2009). Our mixed-effects meta-regression of larval settlement times indicates that reproductive mode and colony morphology traits account for most of the variation in settlement time previously attributed to phylogeny in the random-effects-only model (**Table S4**). This is likely indicative of closely-related species being more likely to share life-history strategies (Darling et al., 2012). Although phylogeny does not explain much variation additional to that explained by life-history traits, our results suggest that phylogenetic relationships could be useful for predicting larval durations in species where these traits are unknown (Macleod et al., 2024). This is encouraging when considering that larval dispersal research has a much greater coverage of coral diversity at the genus-level, especially for Threatened corals (**Figure 1c**). Many biophysical models use larval duration parameters derived from closely related species in the same genus or family (**Figure 3**). Our results validate the effectiveness of this approach for estimating dispersal potential when species-specific information is limited (Macleod et al., 2024). The meta-regressions testing biogeographic variables show that none had a significant effect on either larval settlement or survival times, suggesting that information from corals of similar regions is less useful than life-history traits for inferring dispersal potential in under-studied corals.

Whilst the variability in mean coral larval survival times is significant (**Table 1**), this variability was not significantly driven by life-history traits or phylogenetic relationships. Life-history trait variables did not contribute towards a significant amount of heterogeneity in mean larval survival times (**Table 2**, **Figure 4**). Phylogenetically-controlled random-effects models did not fit the mean larval survival time effect sizes better than equivalent models that did not account for phylogeny (**Table 1**). One explanation for these results could be that differences in experimental conditions, which include both lab-and field-based setups that vary considerably across the literature, altered larval survivorship between studies, confounding any underlying patterns in larval mortality (Connolly and Baird, 2010; Graham et al., 2008; Suzuki et al., 2011). Alternatively, our results may indicate that larval settlement times contribute more than survival times towards observed differences in realised dispersal between coral species with different life-history strategies (van der Ven et al., 2021). Several recent studies have highlighted that larval behavioural responses to settlement cues and the development of settlement competency, both of which contribute to larval settlement time, are important mediators of potential and realised coral larval dispersal (Aoki et al., 2024; Connolly and Baird, 2010; Randall et al., 2024; Strader et al., 2018). Our differing survival and settlement results may suggest that settlement competence and associated behaviours are more predictable and thus more relevant than larval survival for predicting larval dispersal across a variety of data-poor species.

Future work focusing on experimental methods and analysing coral life-history strategies may be needed to overcome some of the limitations in our findings. All mixed-effects meta-regression models had significant residual heterogeneity values (**Table 2**), and mean larval duration values varied greatly within some species (**Figure 4**). This indicates that some larval duration estimates are inconsistent between studies, suggesting a need for further experimental work to assess the full range of mean larval duration values for each species and the extent that larval durations differ between coral life-history strategies (Randall et al., 2024). Traits not included in our meta-analysis could also be important drivers of unexplained variation in coral larval duration (e.g. growth rate). Trait data remains limited or entirely lacking for many coral species. Future research could address gaps in trait data and issues with correlations between traits by conducting ordination analyses to integrate coral life-history trait data together and classify life-history strategies across more coral species (Blomberg and Todorov, 2025; Darling et al., 2012). This would enable more direct examination of potential relationships between coral life history strategies and larval dispersal potential. Studying genera currently not represented in this review would be an effective way to address taxonomic and regional knowledge gaps in coral larval dispersal. Given the time and costs involved in studying realised larval dispersal, measuring simpler, commonly-measured life-history traits and linking these to broader patterns in dispersal potential may be the fastest way to predict connectivity dynamics in under-researched coral populations in the immediate future.

## Conclusion

We assessed variation in reported coral larval durations, finding that life-history traits, specifically reproductive mode and colony morphology, drive a significant amount of variation in mean larval settlement times. This finding indicates that life-history strategies influence larval dispersal potential in corals and suggests that life-history trait data could be used to predict dispersal potential in coral populations lacking larval duration data. We show coral larval settlement times to be phylogenetically non-independent, with the contribution of phylogeny to variation in settlement times being largely explained by reproductive mode and colony morphology, confirming that information from closely-related species is useful for predicting dispersal potential in corals without trait data available. We also find that there are considerable regional and taxonomic knowledge gaps in the coral larval dispersal literature but that spatial patterns do not drive variability in coral larval dispersal potential. Our results suggest that corals differing in reproductive mode and colony morphology could have different requirements for spatial planning resulting from differing larval durations. In the absence of more detailed trait data, future work modelling corals with different reproductive modes and morphologies could predict how realised dispersal varies between coral life-history groupings in specific regions with different oceanographic conditions.

## Acknowledgements

The authors thank Russell Milne for his supervision and for providing valuable feedback during the early stages of this project. The authors also thank Damaris Landers for her advice and expertise during the data collection and analysis stage of this project.

## Author Contributions

All authors conceived the ideas and designed methodology; M. Morecroft, A. Greiner, and L. Faber collected the data; M. Morecroft analysed the data; M. Morecroft led the writing of the manuscript. All authors contributed critically to the drafts and gave final approval for publication.

## Statement on inclusion

Our study was a global review and was based on a meta-analysis of secondary data rather than primary data.

## Data Availability Statement

All data and code used in the systematic review, meta-analysis, and to produce figures are available from the following GitHub repository: https://github.com/matthewmorecroft/coral-larvae-meta and are archived on Zenodo: https://doi.org/10.5281/zenodo.23044993.

