## Supporting Information for "Life-history traits explain variation in the larval dispersal potential of corals"

**Contents**

| **Supporting Methods** |  |  |
| --- | --- | --- |
|  | 1 - Literature search protocols | Page S2 |
|  | 2 - Data extraction protocols | Page S2 |
|  | 3 - Additional life-history trait data | Page S2 |
| **Supporting Figures** |  |  |
| S1 | PRISMA flow-chart | Page S3 |
| S2 | Phylogenies generated for species in meta-analysis | Page S4 |
| **Supporting Tables** |  |  |
| S1 | Sub-categories of studies excluded from systematic review | Page S5 |
| S2 | Categories assigned to each stony coral morphology value | Page S6 |
| S3 | Species with manually assigned morphological categories | Page S6 |
| S4 | σ^2^ values for random-effects variance components | Page S7 |
| S5 | Overall mean settlement times for life-history groupings | Page S7 |
| **References** |  | Page S7 |

**Supporting Methods**

**1 - Literature search protocols**

To identify publications relevant to coral larval dispersal, we developed a 3-part search string, which was as follows: **“(***coral****)** **AND** **(***dispers**-**OR**-*connectivity*-**OR**-*transport**-**OR**-*duration*-**OR**-*distance*-**OR**-*PLD***)** **AND** **(***larv**-**OR**-*propagul**-**OR**-*egg**-**OR**-*sperm**-**OR**-*planul****)**”

To maintain similar search methodology between the two databases and reduce the number of non-relevant studies identified, we applied the search string to the “*Topic*” (titles, abstracts, and indexing) field on the Web of Science database and to the “*Article title*”, “*Abstract”*, and “*Keywords*” fields on the Scopus database. To ensure search reproducibility, we searched records in the Web of Science database published from January 1980 to June 2024 and from January 1980 to January 2024 on the Scopus database (as refining results within a year was not possible for Scopus).

**2 - Data extraction protocols**

Larval survival and settlement data was extracted from the literature using the following protocols. Where a study explicitly stated the mean and SE for larval settlement or survival time, this was recorded (standard deviation, variance, or 95% confidence interval values were converted to SE if necessary). If only raw data on larval survival or settlement over time was reported, these values could be calculated using the frequency of larval deaths or settlements reported for each timepoint. When these frequencies were only reported in plots, the plotted data points were extracted using “WebPlotDigitizer” (Rohatgi, 2024). If neither a mean duration nor sufficient raw data were available, we used the outputs of any models determining mortality or competency loss rates (with confidence intervals) from empirical data. We took the inverse of mean rates and of the upper and lower bounds of the confidence intervals for the rates to calculate mean durations with SE. When none of the above information was available, but the minimum, lower quartile, median, upper quartile, and maximum durations were reported, these values could be used to estimate the mean and SE using the 5-number summary function in the “*metafor*” R package (v4.8-0) (Viechtbauer, 2010).

**3 - Additional life-history trait data**

We used colony morphology data downloaded from the Coral Trait Database in our meta-analysis (IDs of traits used: 183, 627, 104; accessed: 05-05-2025) (Madin et al., 2016). All (except for 6) species in the meta-analysis had data available on their colony morphology. To assign a colony morphology to each coral species included in the meta-analysis, we used the data available under the “*Growth form:* *typical hexacorallia*” trait to classify colony growth forms of stony corals into broader morphological categories (“branching”, “domed”, or “plating”), following the approach of Darling et al. (2012) (**Table S2**). Soft corals were already classified into higher-level morphological categories under the “*Type of growth*” trait (Gómez-Gras et al., 2025). For 6 coral species without morphology information available from the Coral Trait Database, colony morphology was assigned manually by searching the available literature on each species (**Table S3**).

**Supporting Figures**

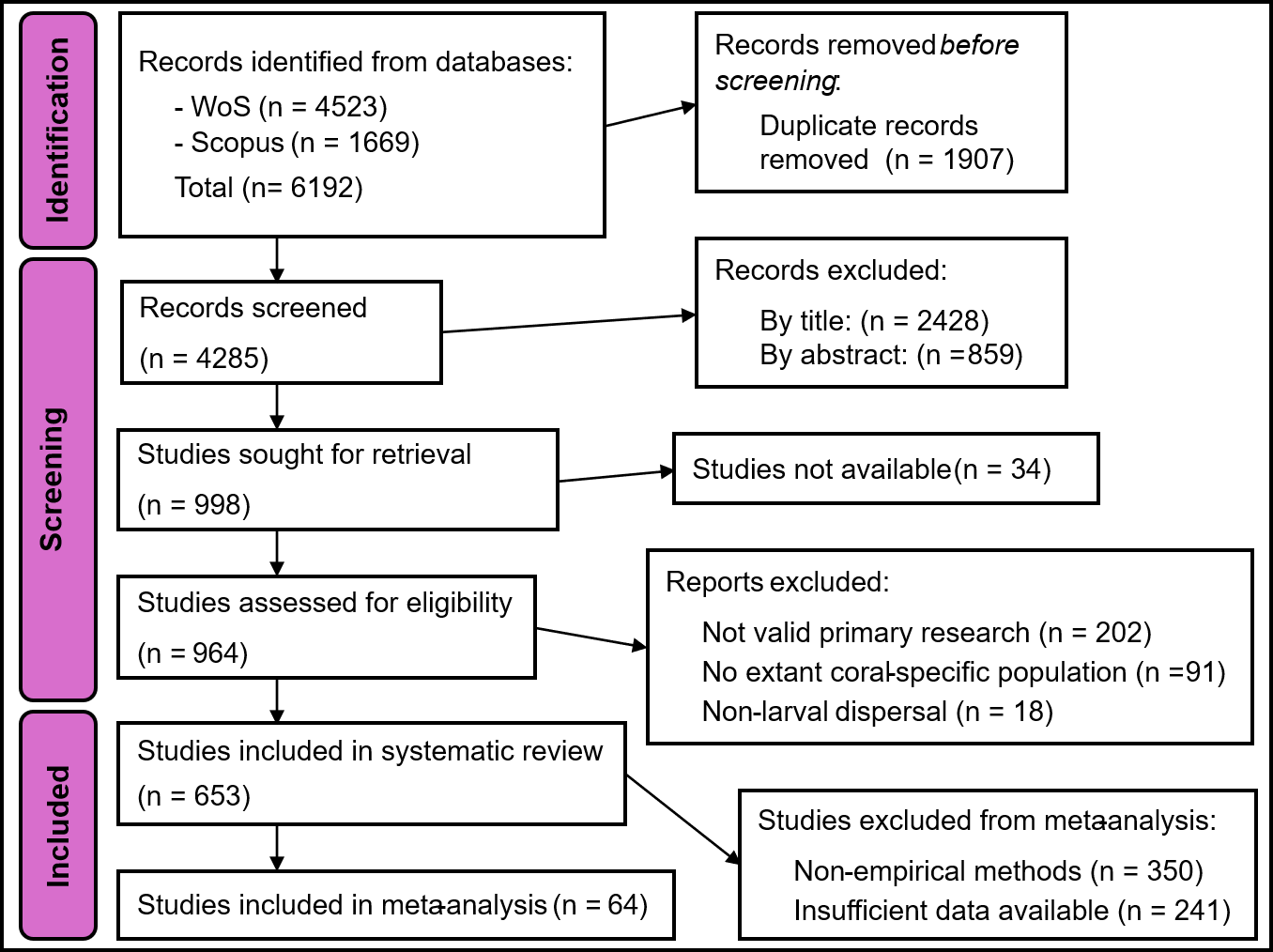

Figure S1: PRISMA diagram (Page et al., 2021) showing work-flow of the literature search and screening carried out in this systematic review of the coral larval dispersal literature, and numbers of studies identified or removed at each stage.

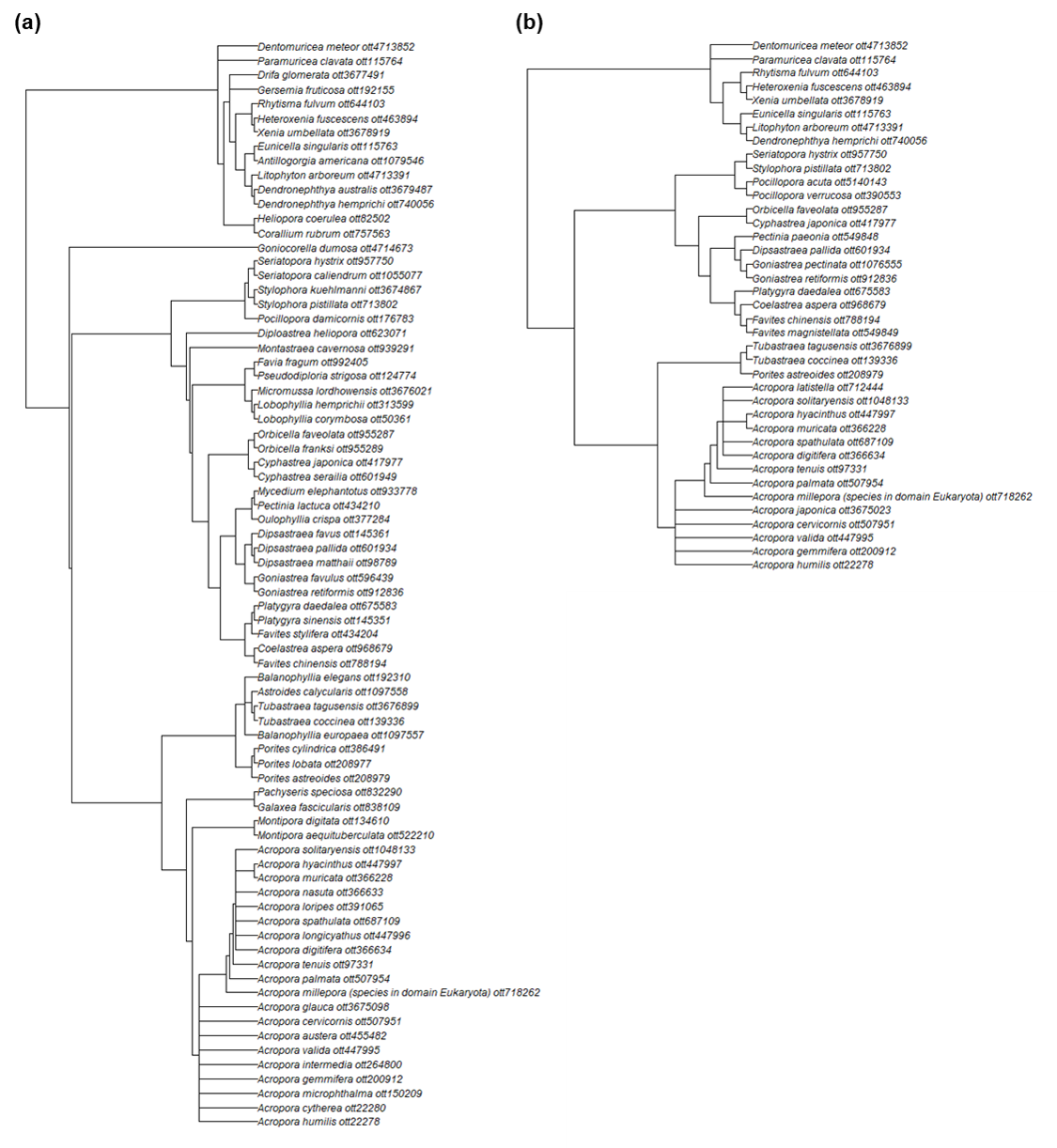

**Figure S2:** Phylogenies obtained for (**a**) species with effect sizes for mean time to larval settlement and (**b**) for species with effect sizes for mean larval survival time.

Table S1: Sub-categories of studies excluded from the systematic review and meta-analysis for failing to meet exclusion criteria 1-3, with a definition of the sub-category and the number of results falling into the sub-category provided.

| Inclusion criteria not met | Sub-category | Sub-category definition | Number of results |
| --- | --- | --- | --- |
| 1 | Commentary | Paper only provides commentary on another study. | 3 |
|  | Grant proposal | Result is a proposal only, does not contain results of research. | 60 |
|  | Retracted | Paper has been retracted. | 1 |
|  | Review | Paper is a review, synthesises information from other studies. | 138 |
| 2 | Generic larvae | Study is of larvae that do not explicitly represent corals. | 50 |
|  | Theoretical | Does not study a real-world existing population. | 18 |
|  | Hydrocoral | Study is of members of the genus Millepora (“fire corals”), which are actually hydrozoans with a distinct motile medusa stage. | 4 |
|  | Palaeontology | Study is of non-extant coral populations. | 19 |
| 3 | Fragmentation | Study is of dispersal via fragmentation, a form of non-larval asexual reproduction. | 1 |
|  | Gametes only | Study is of coral gametes, but only up to the point of fertilisation. | 8 |
|  | Rafting | Study is of dispersal via rafting. | 6 |
|  | Release only | Study is of the release of larvae, but not their post-release dispersal. | 3 |

Table S2: Categories assigned to each value for stony coral morphology from the Coral Trait Database.

| Category | CTD morphology |
| --- | --- |
| Domed | *encrusting* |
|  | *massive* |
|  | *submassive* |
| Branching | *branching_closed* |
|  | *branching_open* |
|  | *hispidose* |
|  | *corymbose* |
|  | *encrusting_long_uprights* |
|  | *digitate* |
| Plating | *laminar* |
|  | *tables_or_plates* |

Table S3: Morphological categories manually assigned to stony coral species without morphology information available on the Coral Trait Database.

| Species | Morphology | Justification |
| --- | --- | --- |
| *Acropora kenti* | branching | Described as “branching” in (Messer et al., 2024) |
| *Astroides calycularis* | domed | General description and images in (Goffredo et al., 2011) match closest to domed category |
| *Balanophyllia elegans* | domed | General description in (Gerrodette, 1981) matches closest to domed category |
| *Goniocorella dumosa* | branching | Described as “branching” in (Beaumont et al., 2023) |
| *Tubastraea coccinea* | domed | General description and images in (Silva et al., 2011) match closest to domed category |
| *Tubastraea tagusensis* | domed | General description and images in (Silva et al., 2011) match closest to domed category |

Table S4: Table summarising the σ^2^ values of the random-effects variance components in the meta-analysis models. Models with “species” as a random effect refer to models which control for phylogeny.

| Model | Random-effects variance component | Effect size: Mean larval settlement time | | |  | Effect size: Mean larval survival time | | |
| --- | --- | --- | --- | --- | --- | --- | --- | --- |
|  |  | σ^2^ | σ | Levels |  | σ^2^ | σ | Levels |
| study \| observation | study | 95.665 | 9.781 | 57 |  | 138.772 | 11.78 | 27 |
|  | observation | 79.438 | 8.913 | 137 |  | 488.001 | 22.091 | 57 |
| study \| observation + species | study | 51.354 | 7.166 | 57 |  | 126.544 | 11.249 | 27 |
|  | observation | 81.369 | 9.021 | 137 |  | 420.252 | 20.5 | 57 |
|  | species | 29.065 | 5.391 | 76 |  | 164.477 | 12.825 | 39 |
| study \| observation + species + morphology* reproductive mode | study | 30.668 | 5.538 | 57 |  | 130.255 | 11.413 | 27 |
|  | observation | 87.891 | 9.375 | 137 |  | 486.297 | 22.052 | 57 |
|  | species | 0.000 | 0.002 | 76 |  | 0.000 | 0.015 | 39 |
| study \| observation + morphology*reproductive mode | study | 30.668 | 5.538 | 57 |  | 130.255 | 11.413 | 27 |
|  | observation | 87.891 | 9.375 | 137 |  | 486.297 | 22.052 | 57 |

Table S5: Table summarising the overall mean larval settlement time for each life-history group predicted by the mixed-effects meta-regression for mean settlement time with colony morphology and reproductive mode as moderator variables. Groups with no data are not included. Data corresponds to vertical lines in Figure 4a.

| Colony morphology | Reproductive mode | Group mean settlement time | SE |
| --- | --- | --- | --- |
| Branching | broadcasting | 13.87 | 1.95 |
|  | brooding | 5.26 | 2.69 |
| Domed | broadcasting | 12.99 | 2.09 |
|  | brooding | 8.74 | 3.52 |
| Plating | broadcasting | 14.09 | 3.69 |
| Erect branched | broadcasting | 27.72 | 5.03 |
|  | brooding | 29.11 | 3.36 |
| Horizontal unbranched | brooding | 9.92 | 5.42 |
